# Assessing flower-visiting arthropod diversity in apple orchards through environmental DNA flower metabarcoding and visual census

**DOI:** 10.1101/2022.01.24.477478

**Authors:** Nerea Gamonal Gomez, Didde Hedegaard Sørensen, Physilia Ying Shi Chua, Lene Sigsgaard

**Affiliations:** Department of Plant and Environmental Science, Section of Organismal Biology, University of Copenhagen, Thorvaldsensvej 40, Frederiksberg C, 1871, Denmark; Section for Evolutionary Genomics, Globe Institute, University of Copenhagen, Øster Farimagsgade 5, 1353 Copenhagen K, Denmark; Wellcome Sanger Institute, Wellcome Trust Genome Campus, Hinxton, Saffron Walden, CB10 1RQ, United Kingdom

**Keywords:** Agricultural systems, biodiversity, environmental DNA, High-throughput sequencing, molecular techniques, insect pollinators

## Abstract

Arthropods are essential to maintaining healthy and productive agricultural systems. Apples are cultivated worldwide and rely on pollination. Honey bees are used for pollination but wild bees and other arthropods also contribute to pollination. Flower visitors can also be natural enemies or herbivores. In some cases, such as Syrphids, a group can have more than one role, adults being pollinators and the larvae being natural enemies of pests. In the present study, we assessed the biodiversity of arthropod flower visitors in four Danish apple orchards and compared the use of molecular and non-molecular techniques to study arthropod communities in agricultural ecosystems. Arthropod DNA collected from apple flowers was analysed by metabarcoding and pollinators were recorded through visual assessment in the orchards. These techniques resulted in two complementary lists of arthropods detected. Non-bee arthropods constituted a big part of the community of apple flower visitors by both methods. Metabarcoding detected 14 taxa and had 72% species resolution while visual census identified 7 different taxa with 14% species resolution. This study showed the importance of using different sampling methodologies to obtain a more accurate picture of fauna present. It also revealed the high presence of non-bee arthropods visiting flowers in apple orchards. The outcome of our study provides information regarding the effects of management practices on arthropod biodiversity, which can contribute to informing on suitable management practices to increase crop yield and maintain healthy agricultural systems.

## Introduction

Arthropod biodiversity in agricultural systems is crucial to maintaining the agroecosystems’ health resilience (Rader et al., 2016; McKerchar et al., 2020; Pardo and Borges, 2020; Angelella et al., 2021). Thus, pollinators and natural enemies of insect pests provide important ecosystem services (Saunders et al., 2016; McCravy, 2018; Porcel et al., 2018; Todd et al., 2020). Many of these species are currently facing multiple threats linked to anthropogenic activities such as land management, pesticides, and invasive species (García and Miñarro, 2014; McKerchar et al., 2020; Pardo and Borges, 2020).

Species diversity, or more specifically complementarity, has been demonstrated to increase crop productivity and sustainability (Klein, 2011; Russo et al., 2015; Rader et al., 2016; McCravy, 2018; Nunes-Silva et al., 2020). The diversity and abundance of pollinators and natural enemies of insect pests are limited by the amount of non-crop flowering habitat (hedgerows or flower strips) and pest management practices (Rader et al., 2020; Hambäck et al., 2021). Therefore, the conservation of semi-natural habitats in managed landscapes, such as apple orchards, is crucial to improving arthropod services (Klein et al., 2007; Joshi et al., 2016; Herz et al., 2019; Pardo and Borges, 2020).

Apple is a perennial crop with 1500 ha cultivated in 2020 in Denmark (EU Commission, 2021). It is a highly pollinator-dependent crop, and most studies of pollinators have been focused on Hymenopteran species (Földesi et al., 2020; Pardo and Borges, 2020; Weekers et al., 2022). However, apple fauna includes a broad diversity of arthropod pollinators like Coleoptera, Diptera and, Lepidoptera, of which some are also natural enemies of insect pests (Rader et al., 2016; Lucas et al., 2018; Herz et al., 2019; Földesi et al., 2020).

Arthropods are influenced by climatic and meteorological conditions, exhibit year-to-year and day-to-day variation, and can be small, fast and cryptic (Gibbs et al., 2017; Lucas et al., 2018). For this reason, a combination of methods will provide a more complete overview of arthropods present in apple orchards. Traditional techniques such as visual assessment rely on the observant’s knowledge and ability to capture and identify arthropod species, and the methods can relatively be labour-intensive (Russo et al., 2015; Evans and Kitson, 2020). Visual assessment is attractive in providing direct observation data of flower visiting arthropods, but observations require weather conditions that allows pollinator activity, which can be limited in Denmark in spring.

High-throughput sequencing (HTS) technologies have recently been applied in the study of arthropod communities in managed ecosystems. One such approach, metabarcoding, consists of the amplification and detection of a specific sequence of interest from a mixture of environmental DNA (Ruppert et al., 2019; Evans and Kitson, 2020). This technique has many applications, including pollen identification from different sources such as plant tissues or animal traces (Lucas et al., 2018; Evans and Kitson, 2020). Metabarcoding of remnant arthropod DNA on plant samples is a relatively new tool that has just started to be used for plant-arthropod network research (Ruppert et al., 2019; Thomsen and Sigsgaard, 2019). Compared to traditional methods, metabarcoding is faster in the field, and can potentially provide a more reliable overview of apple flower visitors (Ruppert et al., 2019; Evans and Kitson, 2020), with better taxonomic resolutions, while being more cost-efficient (Yang et al., 2014; Chua et al., 2021). However, degraded DNA and unsuitable sample processing might lead to skewed results (De Barba et al., 2014; Deagle et al., 2019). Moreover, this methodology is not appropriate for species abundance quantification studies (Lamb et al., 2019; Evans and Kitson, 2020; Todd et al., 2020).

We assessed the diversity of flower-visiting arthropods in four Danish apple orchards using metabarcoding of environmental DNA (eDNA) from apple flowers and visual assessment of all flower visitors present in study sites. Through the combination of these two techniques, we aimed to create a more complete list of arthropod flower-visitors to apple orchards and study their distribution in and across the orchards. Based on existing knowledge of the apple orchard arthropod community and floral visitors (Cross et al., 2015; Alford, 2019; Cahenzli et al., 2019; Thomsen and Sigsgaard, 2019), we predict that i) pollinator diversity will be mainly represented by Hymenoptera and Diptera, ii) there will be significant differences in arthropod communities across the orchards, and iii) arthropods detected will be similar between the methodologies used.

## Methods

Arthropods were sampled in four apple orchards on Sealand, Denmark; one 20 km north of Copenhagen (Frydenlund), two 25 km and 37 km south-west of Copenhagen respectively (Kildebrønde, and Ventegodtgaard), and finally the Pometum, 16 km west of Copenhagen, belonging to University of Copenhagen (Fig. 1A and see Supplemental information 1 Table S1). Sites were separated by at least 9 km and located in an agricultural matrix. While Frydenlund and Kildebrønde were relatively large orchards with more than 20 rows of apples (more than 100m wide), apple plots at the Pometum and Ventegodtgaard were only 7 and 10 rows wide respectively (less than 40m wide). Ventegodtgaard was managed organically while the other three followed integrated pest management (IPM). All the orchards had honey bee hives within the field (Frydenlund and Kildebrønde) or in the surrounding crops (the Pometum and Ventegodtgaard). Arthropods were sampled in four different distances from the margin (side of orchard where flowers strip was sown) of the orchard: row 1 (0 m from the margin, edge of the orchard (first row of apple trees)), row 3, row 5 and row 10, approximately 5, 10 and 25m from the margin, respectively (Fig. 1B). In the smallest orchard, at the Pometum, the last row was row 7, approximately 15m inside and facing a strip of grass followed by a pear orchard (Fig. 1B). The visual assessment protocol was adapted from *Westphal* et al. (2008). The vegetation surrounding the orchards was different across them (See Supplemental information 1 Table S2). Sampling was conducted when the percentage of open apple flower buds was between 50-90%. This occurred in late May 2020 for two weeks till the end of the apple flowering period.

**Figure 1.**
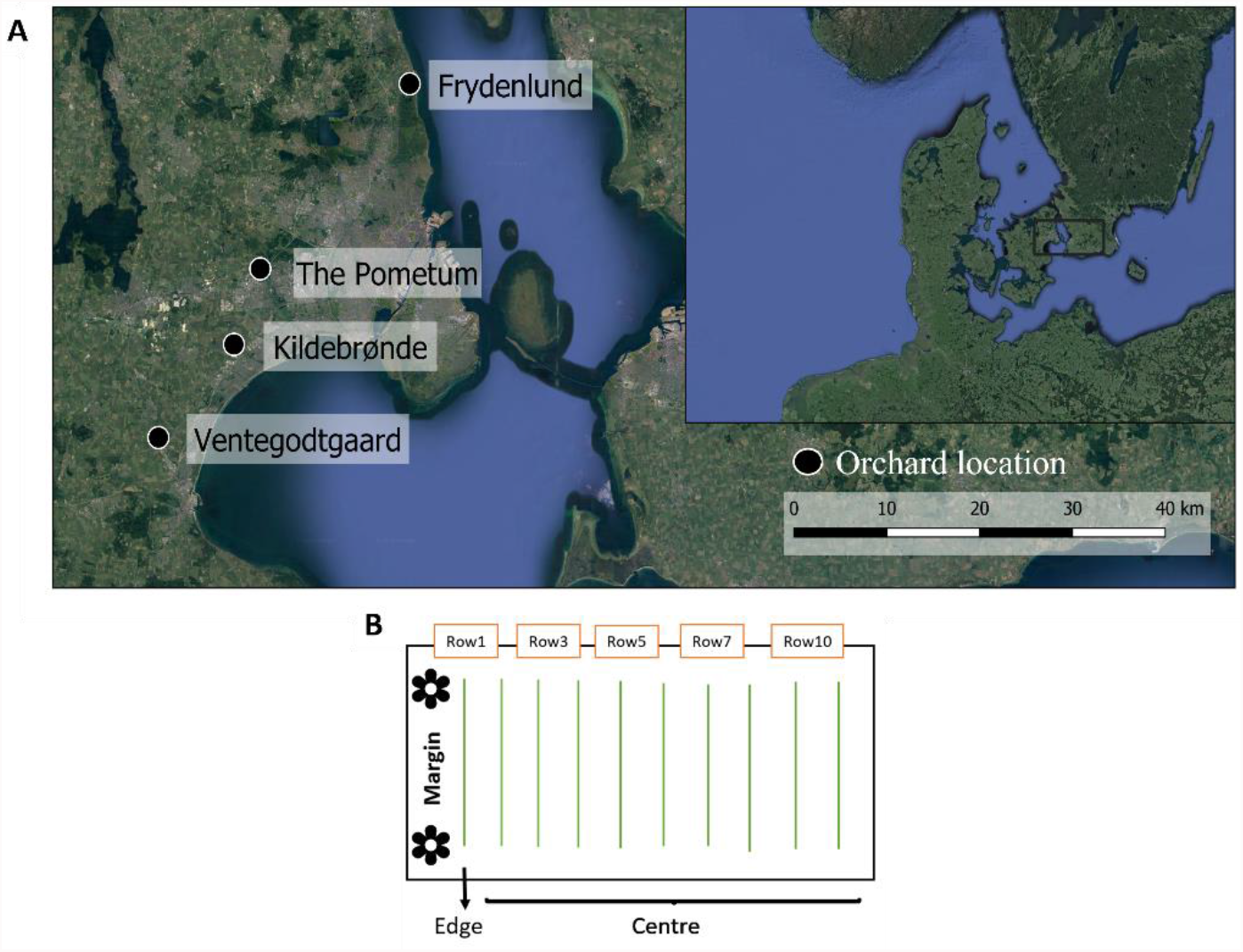
**A** Location of the four apple orchards where apple flowers (n=60) and arthropods (n=279) were collected and **B** rows disposition at each orchard.

### Metabarcoding

We followed the methods from Thomsen and Sigsgaard (2019) to analyse the environmental DNA (eDNA) present in the sampled apple flowers.

In the morning, five individual apple flowers were collected in rows 1, 3 and 7 of each orchard. Thus, a total of 60 flowers were individually collected in sterile plastic tubes (50mL, Thermo Scientific). Collection was done using single-use sterile nitrile gloves, to avoid any pollution of samples. Plastic tubes were put on ice and stored at −20ºC prior to DNA extraction.

### DNA extraction

DNA extraction was carried out at the Department of Plant and Environmental Science laboratories, University of Copenhagen. The experiment was performed in a PCR-free laboratory to prevent contamination. DNA was extracted using the Qiagen DNeasy Blood & Tissue Kit and protocol in a flow hood. First, the whole apple flower was transferred to a 2ml Eppendorf tube prior to DNA extraction. Lysis was performed by adding 900μl of a cell lysis solution (ATL buffer (QIAGEN)) and 100 μl of proteinase K (QIAGEN). Samples were disrupted in the TissueLyser II (QIAGEN) for 2 minutes at 30Hz and incubated at 56ºC with agitation in a rotor for 3 hours. Samples were vortexed for 10 seconds before transferring 800 μl of the lysis mixture to a new 2ml Eppendorf tube. 800 μl of lysis buffer (AL buffer (QIAGEN)) was added and the mixture was mixed thoroughly by vortexing before incubation at 56ºC for 10 minutes. 800 μl of absolute ethanol was added to the mixture, followed by vortexing before adding the mixture to the spin columns. The mixture was spun through the membrane filter over three rounds (700 μl per round) with 1.5 minutes of centrifugation at 8000rpm after each round. The flow-through was discarded every round. The Spin columns were washed by adding 600 μl of wash buffer (AW1 (QIAGEN)) and centrifuged for 1.5 minutes at 8000rpm, followed by adding 600 μl of AW2 (QIAGEN) and centrifuged for 3.5 minutes at 14000rpm. Each spin column was transferred to a new 2ml Safe-lock Eppendorf tube and DNA was eluted in 2×60 μl AE buffer (QIAGEN) with a 15 minute incubation step at 37ºC before centrifugation (1.5 minutes at 10000rpm). One extraction blank was included at the beginning of the process to test for possible contamination during the procedure.

DNA extracts from apple flowers collected from the same row, but different apple trees in the same orchard, were pooled according to the DNA concentration and 260/280 ratio, measured by Microvolume spectrophotometer (mySPEC, VWR) (See Supplemental information 1 Table S3). This resulted in a total of 36 pooled DNA extracts and one extraction blank stored at −20ºC prior to further analysis.

### PCR amplification

All DNA extracts including the extraction blank were sent to *AllGenetics & Biology SL* (www.allgenetics.eu) for PCR amplification and sequencing. For eDNA flower metabarcoding, a 157 bp fragment of the COI genomic region was amplified using the primers ZBJ-ArtF1c (5’ AGA TAT TGG AAC WTT ATA TTT TAT TTT TGG 3’) and ZBJ-ArtR2c (5’ WAC TAA TCA ATT WCC AAA TCC 3’) (Zeale et al., 2011; Thomsen and Sigsgaard, 2019). Three PCR replicates were generated for each sample.

PCR reactions for each sample were carried out in a final volume of 12.5 μL, containing 1.25 μL of template DNA, 0.25 μM of the primers, 6.25 μL of Supreme NZYTaq 2x Green Master Mix (NZYTech), CES 1x, and ultrapure water up to 12.5 μL. The PCR reaction was incubated as follows: an initial denaturation step at 95 ºC for 5 minutes, followed by 35 cycles of denaturing at 95 ºC for 30 seconds, annealing at 49 ºC for 45 seconds, 72 ºC for 45 seconds, and a final extension step at 72 ºC for 7 minutes. The oligonucleotide indices which are required for multiplexing different libraries in the same sequencing pool were attached in a second PCR round with identical conditions but using only five PCR cycles and 60 ºC as the annealing temperature. A PCR blank that contained 1.25 μL of ultrapure water instead of DNA (BPCR) was included to check for contamination during PCR library preparation. Additionally, PhiX Control v3 (Illumina) was used as a control library for Illumina sequencing runs.

The PCR libraries were run on 2 % agarose gels stained with GreenSafe (NZYTech) and imaged under UV light to verify the library size (227pb). PCR libraries were purified using the Mag-Bind RXNPure Plus magnetic beads (Omega Biotek), following the instructions provided by the manufacturer. PCR libraries were pooled together in equimolar amounts. Pooled PCR libraries were purified using the Mag-Bind RXNPure Plus magnetic beads (Omega Biotek). The pooled PCR libraries were sequenced in 3/16 of an Illumina MiSeq PE300 run.

### Data analysis

Illumina paired-end raw files consist of forward (R1) and reverse (R2) reads sorted by PCR libraries and quality scores. The indices and sequencing Illumina adapters were trimmed during the demultiplexing step. Any remaining Illumina adapters were removed using the software CUTDAPT (Martin, 2011). The resulting trimmed sequences were used for further analysis.

The analysis of the trimmed sequences was carried out using the OBITools package, which allows sorting and filtering of sequences based on the taxonomy (Boyer et al., 2016). A total of 36 samples, one extraction blank (sample 37), and one PCR blank (BPCR) were included in the dataset. Forward and reverse reads were first merged (ILLUMINAPAIREDEND) and unaligned sequence records were removed with an alignment score below 40 (OBIGREP). Reads were dereplicated into unique sequences by OBIUNIQ. Sequences with only a single copy (singletons) and shorter than 100 bp were removed by OBIGREP. Amplification and sequencing errors generated during PCR and sequencing were identified and cleaned with OBICLEAN, using a threshold ratio of 5% (De Barba et al., 2014).

For Taxonomic assignment, the NCBI reference database (Deiner et al., 2017) was built through ecoPCR. Taxonomic assignment for Zeale primers (Zeale et al., 2011) was performed using the ECOTAG program, which compares each sequence of the data set to the created taxonomic database (Boyer et al., 2016). Post-OBITools sequence filtering and merging of the taxonomic assignments were carried out with R-studio 1.4.1103 (R Core Team, 2020). Following the analysis detailed in Chua et al. (2021), only sequences that fulfilled the following criteria were kept: i) matched 98% of our reference database, ii) had a minimum of three reads of each taxon observed within each PCR replicate (See Supplemental information 1 Table S4), and iii) occurring in at least two PCR replicates of a sample (Ficetola et al., 2015; Rasmussen et al., 2021b) (See Supplemental information 1 Table S5). No sequences were found in the extraction blank and PCR blank when checked for possible contamination. The remaining sequences whose taxa identification was higher than species level were compared to BLAST, changing the final taxa name from only those with 100% of Query cover and percent identity (See Supplemental information 1 Table S6). Only taxa that belonged to the arthropod phylum were kept (See Supplemental information 1 Table S7). The resulting output was grouped according to four different orders: Blattodea (BT), Coleoptera (CP), Diptera (DI), and Lepidoptera (LP) (See Supplemental information 1 Table S8).

### Visual census

The data from the visual assessment was generated as part of the Beespoke project, where the effect of arthropod flower visitor diversity and abundance was assessed as a function of distance from a flower strip. An observer noted all the arthropod flower visitors 2.5 meters on each side of the observer (covering two rows of apple trees). The type and number of arthropods were recorded for five minutes in each observational transect walk. Arthropods were identified to morpho-groups: Bumblebee (BB), Coleoptera (CP), Diptera (DI), honey bee (HB), Lepidoptera (LP), Syrphid (SY), and wild bee (WB). This classification was established following previous studies on typical pollinators found in apple orchards (Ramírez and Davenport, 2013). We did not sort Hymenoptera to family level but instead sorted them according to honey bees, wild bees and bumblebees as foraging habits and activity differ within the same family (Delaplane et al., 2000; Gardner and Ascher, 2006; Földesi et al., 2020; Pardo and Borges, 2020).

Transects were performed at each orchard during two different times of the day; mornings (between 9:00 and 12:00) and afternoons (between 13:00 and 16:00). We sampled twice in Kildebrønde, the Pometum and Ventegodtgaard, once in the morning and once in the afternoon. We sampled thrice in Frydenlund, once in the morning and twice in the afternoon. Surveys were only carried out when wind speed was not exceeding 7m/sec and a threshold temperature of 10ºC on sunny days, and 15ºC on overcast days (Ramírez and Davenport, 2013). These factors decided when to select the days and times for sampling (using regional weather forecast from Danish Meteorological Institute).

### Statistical analysis

For metabarcoding, statistical analysis was carried out using the R package *vegan* (Oksanen et al., 2020). Only taxonomic units assigned to Arthropoda and present in Denmark were considered (See Supplemental information 1 Table S8). Sequence counts were analysed in two different ways; presence/absence of each insect taxa measured using the frequency of occurrence (Fo), and relative read abundance (RRA) obtained by proportional summaries of counts (Deagle et al., 2019). Read counts were transformed into RRA data using the *decostand* function from R package *vegan* (Oksanen et al., 2020). Summaries based on occurrence data (Fo) are more sensitive to rare diet taxa and pooled samples than RRA (Deagle et al., 2019). Thus, results were discussed using both RRA and Fo values. We used order as a taxonomic unit for metabarcoding diversity analysis to compare results from both methodologies.

For visual assessment, the sampling effort was different across the orchards. Due to meteorological conditions and orchard location, the four orchards were sampled on a different number of days. Honey bee (HB) data was not included in the arthropod richness analysis as it occurs in Denmark only as a managed species (Rasmussen et al., 2021a). We considered morpho-groups as taxonomic units for arthropod composition analysis. The effect of honey bee hives on wild pollinators was studied. Orchards with honey bee hives within the field were considered orchards with close honey bee hives (Kildebrønde and Frydenlund) while those with hives in the surrounding crops were established as orchards with distant hives (Ventegodtgaard and the Pometum). Wild pollinators included all arthropods that were not honey bees. Chi-squared test was used to test for dependency between the variables while the type of relationship was measured by an ODDS ratio and Risk estimate from R package *fmsb* (Nakazawa, 2021).

To avoid biased results due to the small sample size and sampling effort, rarefaction tests were carried out for both the molecular and non-molecular analyses to assess sampling completeness and the relationship between arthropod richness and type of orchard (Gotelli and Chao, 2013; Russo et al., 2015). We used arthropod order and the orchard data to complement and compare both molecular and non-molecular methodologies. Arthropod communities were compared using 95% confidence intervals (Chao1 estimator) of the rarefaction curves and extrapolation of Hill numbers (species richness (q=0)). Differences across the expected diversities are significant when 95% confidence intervals overlap (Chao et al., 2014). The R package *iNEXT (Chao et al., 2014; Hsieh et al., 2016)* from R version 4.0.3, based on the bootstrap method, was used to assess the uncertainty of the proposed sample completeness measure. Orchard (Frydenlund, Kildebrønde, the Pometum, and Ventegodtgaard) was chosen as a sampling effort unit. For the molecular analysis, only occurrence data (Fo) was used as it represented the presence and absence data of the different genera in the orchards.

## Results

### Metabarcoding

From our metabarcoding analyses, 4918 sequences were removed due to either unsuccessful PCR amplification or filtering parameters (See Supplemental information 1 Table S4). The final dataset consisted of 355 sequences with a minimum percentage identity of 98% to at least one taxon in the EMBL reference database after taxonomic assignment (See Supplemental information 1 Table S5). 129 of these sequences belonged to an oomycete (Peronosporaceae family). After merging the sample replicates, the final dataset consisted of 14 insect taxa (See Supplemental information 1 Table S8). The taxonomic resolution was 7% (1 taxa) at order level, 21% at genus level (3 taxa), and 72% at species level (10 taxa) (Table 1). Two detected species (the Diptera *Lonchoptera uniseta* and the Blattodea *Periplaneta americana*) are not naturally occurring in Denmark (DanBIF Secretariat, 2021), except, in the case of *P. americana* as an occasional pest in bakeries (Jensen, 1993). Neither of them had 100% match to BLAST reference dataset. Therefore, *Lonchoptera uniseta* was kept at genus level *(Lonchoptera*) since other species of this genus, such as *Lonchoptera lutea*, are common in the area. It is possible that another species of cockroach may be found in the orchards, most likely *Ectobius lapponicus*, a forest cockroach that belongs to the same family and is common in Denmark (DanBIF Secretariat, 2021). Thus, we proposed this species to the list in Table 1.

**Table 1.** Arthropod composition found across all orchards through flower metabarcoding with the COI Zeale primers. (*) Species that have not been documented in Danish apple orchards. Taxon assignment: 1= final taxon stays the same, 2= final taxon changed to another species in the same order indicated by (), 3= final taxon moved up a taxonomic level due to species not found in Denmark; changes based on Naturbasen (2021) and DanBIF Secretariat (2021). Order: B=Blattodea, C=Coleoptera, D=Diptera and L=Lepidoptera. Resolution final taxa: final level at which taxa is assigned to S=species or G=genus. Life stage according to time of year sampling was done: A=Adult, L=Larva. Functional group: role of arthropod in apple orchards during the specific life stage. Contribution to pollination: 0= none (they do not act as a pollinator), 1= some (they act as pollinators or based on data from related species, have the potential to act as pollinators). References in table footnote

| Taxon assignment | Order | Family name | Genus name | Final taxa | Resolution final taxa | Stage | Functional group | Contribution to pollination |
| --- | --- | --- | --- | --- | --- | --- | --- | --- |
| 2 | B | Blattodea |  | *(Ectobius lapponicus)* | S | A | Omnivore | 1<sup>1</sup> |
| 1 | C | Coccinellidae | *Harmonia* | *Harmonia axyridis* | S | A | Predator | 1<sup>2,3</sup> |
| 1 | C | Nitidulidae | *Brassicogethes* | *Brassicogethes aeneus* | S | A | Herbivore | 0<sup>4</sup> |
| 1 | D | Anthomyiidae | *Delia* | *Delia platura* | S | A | Herbivore | 1<sup>5</sup> |
| 1 | D | Drosophilidae | *Scaptomyza* | *Scaptomyza* sp. | G | A | Herbivore | 1<sup>6</sup> |
| 3 | D | Lonchopteridae | *Lonchoptera* | *Lonchoptera* sp. | G | A | Herbivore, adults feed on nectar | 1<sup>7</sup> |
| 1 | D | Muscidae | *Musca* | *Musca domestica* | S | A | Scavenger, Coprophage | 1<sup>8</sup> |
| 1 | L | Geometridae | *Operophtera* | *Operophtera*<br>*brumata* | S | L | Herbivore | 0<sup>9</sup> |
| 1 | L | Geometridae | *Operophtera* | *Operophtera*<br>*fagata* | S | L | Herbivore | 0<sup>9</sup> |
| 1 | L | Geometridae | *Operophtera* | *Operophtera* | G | L | Herbivore | 0<sup>9,10</sup> |
| 1 | L | Tortricidae | *Pandemis* | *Pandemis*<br>*cerasana* | S | L | Herbivore | 0<sup>10</sup> |
| 1 | L | Yponomeutidae | *Yponomeuta* | *Yponomeuta*<br>*malinellus* | S | L | Herbivore | 0<sup>11</sup> |
| 1 | L | Yponomeutidae | *Yponomeuta* | *Yponomeuta*<br>*cagnagellus* | S | L | Herbivore | 0<sup>12</sup> |
| 1 | L | Yponomeutidae | *Yponomeuta* | *Yponomeuta* | G | L | Herbivore | 0<sup>11,12</sup> |

Footnote: <sup>1</sup>Macgregor and Scott-Brown (2020), <sup>2</sup>Bertrand et al. (2019), <sup>3</sup>Steenberg and Harding (2009), <sup>4</sup>Sutter and Albrecht (2016), <sup>5</sup>Cook et al. (2020), <sup>6</sup>Gibson et al. (2006), <sup>7</sup>Stalker (1956), <sup>8</sup>Ramírez and Davenport (2013), <sup>9</sup>Holliday (1977), <sup>10</sup>Svensson et al. (1999), <sup>11</sup>CFIA (2006), <sup>12</sup>Turner et al. (2010)

### Arthropod composition

The composition of arthropods found in the apple orchards differed based on the metabarcoding analysis used; frequency of occurrence (Fo) and relative read abundance (RRA) (See Supplemental information 1 Table S9). Based on Fo analysis, the occurrence of Diptera (DI) in apple orchards was the highest as compared to the other orders at 44% (Fig. 2A). Whereas, based on RRA analysis, Lepidoptera (LP) had the highest read counts at 49% (Fig. 2B). The least common arthropod groups were Blattodea (BT) and Coleoptera (CP) in both analyses (Fig. 2A, B).

**Figure 2.**
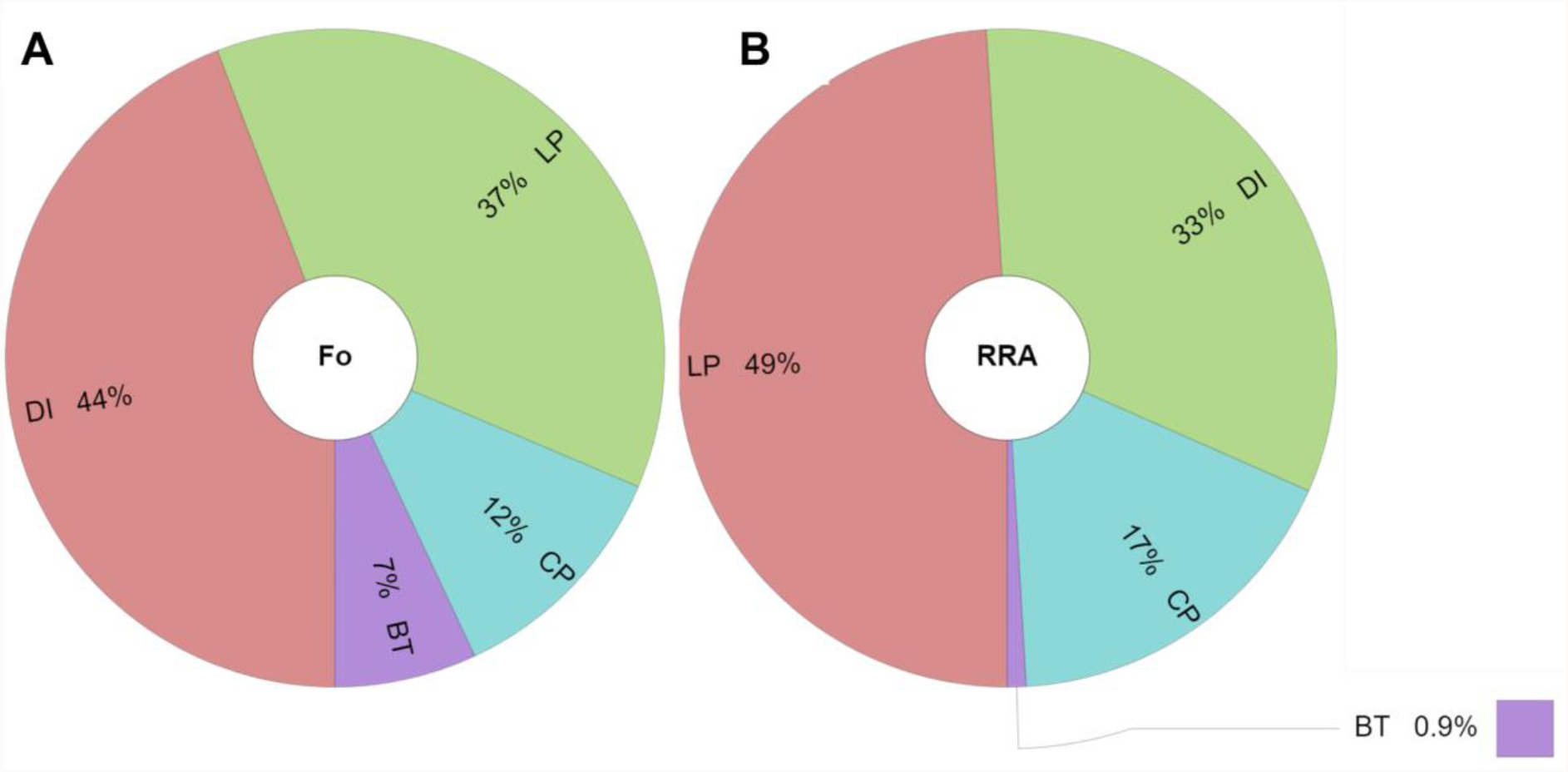
Krona chart showing the percentage of four different arthropod orders identified from COI metabarcoding of flower samples collected in the four orchards based on **A** frequency of occurrence (Fo) and **B** relative read abundance (RRA). Order: Blattodea (BT), Coleoptera (CP), Diptera (DI), and Lepidoptera (LP).

### Arthropod richness

Sampling coverage (SC) indicated sample adequacy at Frydenlund, Kildebrønde and the Pometum, but with values below the estimated total (SC>0.95) in Ventegodtgaard (See Supplemental information 1 Table S10). Thus, estimated values were used to compare arthropod richness (q=0) across sites. Rarefaction curves showed that there were significant differences in arthropod richness (non-superimposed confidence intervals) between Frydenlund (Sestimated=2) and Ventegodtgaard (Sestimated=4.47) (See Supplemental information 2 Fig. S1). All arthropod orders were detected in Ventegodtgaard. Blattodea and Coleoptera were absent from Frydenlund, Blattodea was also absent from Kildebrønde, and Diptera was absent from the Pometum (See Supplemental information 1 Table S9).

### Visual census

#### Arthropod composition

A total of 279 individuals within 7 taxa were identified through visual census. The taxonomic resolution was 43% (3 taxa) at order level, 14% at genus level (1 taxa), 14% (1 taxa) at family level, and 14% at species level (1 taxa). The remaining taxa (14%) were identified to higher taxonomic levels. The most abundant arthropod was honey bee (47%) followed by Diptera (38%) (Fig. 3). The least abundant arthropods detected were bumblebees and Syrphidae (5% each), Coleoptera (3%), wild bee (2%), and Lepidoptera (0.7%). Non-bee pollinators constituted 47% of total individuals recorded.

**Figure 3.**
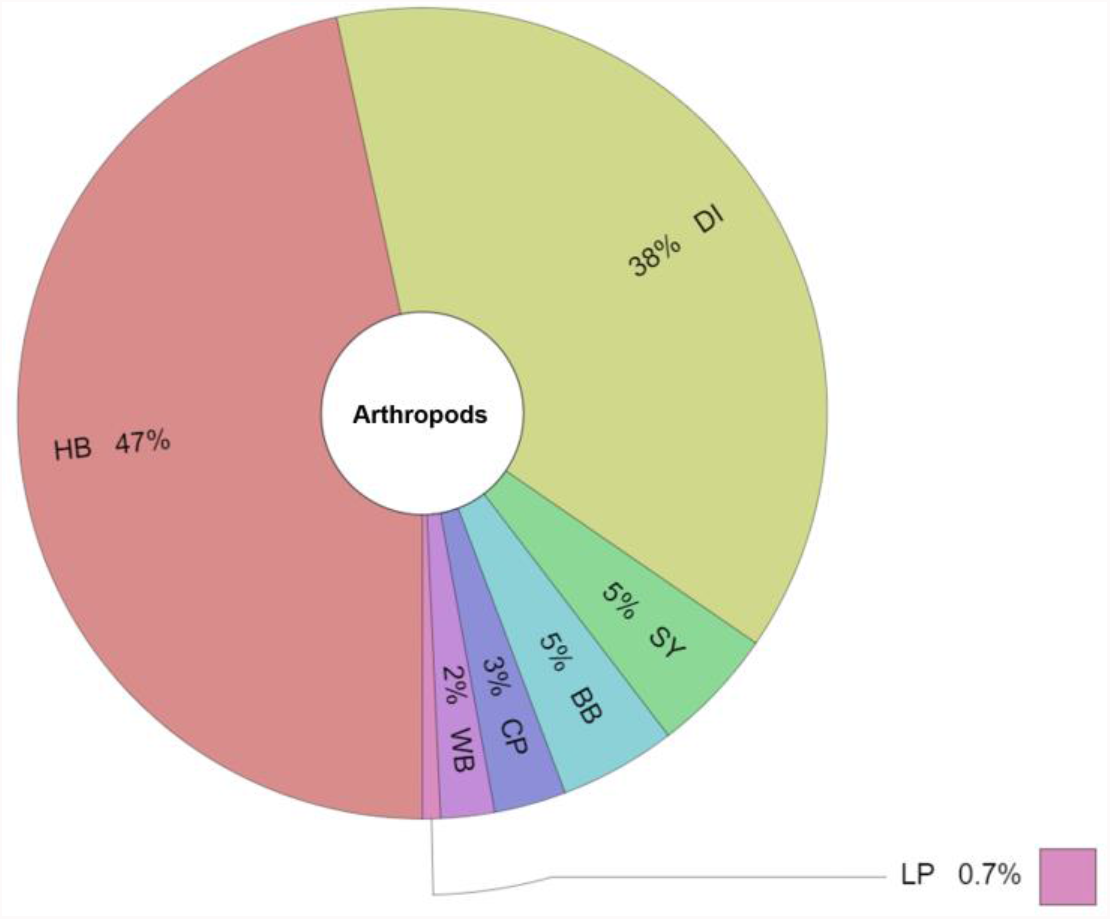
Krona chart showing the percentage of arthropod morpho-groups recorded by visual census in the four Danish apple orchards. Arthropods: Bumblebee (BB), Coleoptera (CP), Diptera (DI), honey bee (HB), Lepidoptera (LP), Syrphid (SY), and wild bee (WB).

#### Arthropod richness

Sampling coverage (SC) indicated sample adequacy at Frydenlund and Ventegodtgaard, but with values below the estimated total (SC>0.95) in Kildebrønde and the Pometum (See Supplemental information 1 Table S12). Thus, estimated values were used to compare arthropod richness (q=0) across sites. Rarefaction curves showed that there were no significant differences in arthropod richness between the orchards (See Supplemental information 2 Fig S2). All arthropod morpho-groups were detected in Ventegodtgaard. Lepidoptera was absent from Frydenlund, Syrphidae and Lepidoptera from the Pometum orchard, and Coleoptera from Kildebrønde (See Supplemental information 1 Table S11). Regarding the arthropod abundance in the orchards, we detected a lower number of individuals in the Pometum (n=42) as compared to the other orchards (Kildebrønde n=85, Frydenlund n=84, Ventegodtgaard n=68) (See Supplemental information 1 Table S11). However, this changes when we remove honey bees from the analysis. Less wild pollinators were recorded in Kildebrønde and Frydenlund when compared to the Pometum and Ventegodtgaard (See Supplemental information 1 Table S13). Honey bees were more common in orchards with close hives compared with orchards with distant hives (Odds ratio estimate, *CI*=2.63-7.56, *p* =1.193e-08, *estimate*= 4.46), where wild pollinators were observed twice as often as honeybees (Risk ratio estimate, *CI*=1.43-2.13, *p*=1.193e-08, *estimate*=1.75).

#### Comparison between metabarcoding and visual census

Using metabarcoding, we did not detect any Hymenoptera or Syrphidae. However, we detected other orders such as Blattodea (BT) that were not found in the visual census. The arthropods detected from visual census were all adult pollinators that visited apple flowers (Fig. 3). Coleoptera, Diptera, and Lepidoptera were detected using both metabarcoding and visual census (Table 2). Ventegodtgaard was the only orchard where all the three orders were detected using both methods. Arthropod presence differed between orchards (Frydenlund, Kildebrønde, the Pometum and Ventegodtgaard) and methodologies (metabarcoding and visual census) in terms of the orders detected (Table 2). Ventegodtgaard had the highest arthropod richness for both methodologies. Based on metabarcoding, arthropod richness in Frydenlund was the lowest as compared to the other orchards. Whereas, based on visual census, the Pometum had the lowest arthropod richness.

**Table 2.** Arthropod composition found in the four Danish orchards with metabarcoding (frequency of occurrence (Fo)) and visual census. Arthropod richness is shown by the Sestimated by rarefaction analysis results from both methodologies.

|  | Frydenlund |  | Kildebrønde |  | The Pometum |  | Ventegodtgaard |  |
| --- | --- | --- | --- | --- | --- | --- | --- | --- |
| Arthropod | *Fo* | *Visual* | *Fo* | *Visual* | *Fo* | *Visual* | *Fo* | *Visual* |
| Diptera (DI) | 4 | 41 | 8 | 12 | 0 | 27 | 7 | 26 |
| Coleoptera (CP) | 0 | 1 | 1 | 0 | 3 | 1 | 1 | 6 |
| Lepidoptera (LP) | 4 | 0 | 3 | 1 | 4 | 0 | 5 | 1 |
| Arthropod richness (Sestimated) | 2 | 5.98 | 3 | 6.9 | 3 | 4.97 | 4.47 | 6.88 |

## Discussion

We demonstrated in our study the reproducibility of using environmental DNA (eDNA) from flowers to detect arthropod visitors in apple orchards (Thomsen and Sigsgaard, 2019). We further showed the complementarity of using visual census and metabarcoding methodologies to improve current knowledge of arthropod communities regardless of differences in emergence dates or foraging periods. Visual census found that pollinator communities were mainly represented by Hymenoptera and Diptera. Metabarcoding found DNA traces of several herbivores, some of them apples pests, and also revealed other flower-visitors which may be more night-active and hence missed by visual census. The fact that flowers were collected in the morning could explain that DNA traces from day active Hymenoptera were not captured by the molecular technique. Therefore, this study also highlights some challenges to consider when using either techniques for future studies.

### Arthropods identified in apple orchards

From our metabarcoding results, we identified some common apple flower visitors such as Diptera and Coleoptera. Additionally, we detected Blattodea that has not commonly been reported from apple orchards (García and Miñarro, 2014; Rader et al., 2020; Hambäck et al., 2021). These differences might be driven by the detection methods utilised, as traditional assessments used in other orchard arthropod assessment studies might miss cryptic or nocturnal fauna (Russo et al., 2015; Deiner et al., 2017; Evans and Kitson, 2020; Geppert et al., 2020). A broader diversity of arthropods, including predators and parasitoids, were found in one other similar flower metabarcoding study (Thomsen and Sigsgaard, 2019). The lower diversity of arthropods detected in our study (14 taxa) is likely due to our sampling in an agricultural landscape and the sampling period limited to the flowering of apple trees, as well as to differences in experimental settings between studies (Leonhardt et al., 2013; Marquina et al., 2019), where i) their sampling season was in August when higher temperatures are reached and therefore more arthropod visitors are present compared to spring (Ramírez and Davenport, 2013; Rader et al., 2016; Gibbs et al., 2017), ii) they collected seven flower species which would have attracted more flower visitors instead of just one flower collected in our study (apple flowers) (Quinet et al., 2016; Samnegård et al., 2019; Pardo and Borges, 2020), and iii) the use of two primers in their research instead of one (Mitochondrial COI (Zeale et al., 2011) and ribosomal 16S (Elbrecht et al., 2016)). This could also be the reason why we did not recover any sequences belonging to Hymenoptera and Syrphidae, as the COI primer that we used is a generic primer and could have failed to amplify DNA from these taxa (Lamb et al., 2019; Marquina et al., 2019; Rasmussen et al., 2021b). The Zeale primer also non-selectively amplifies DNA fragments from other taxonomic groups other than arthropods (De Barba et al., 2014; Thines and Choi, 2016; Lamb et al., 2019; Marquina et al., 2019; Rasmussen et al., 2021b), which is demonstrated by the presence of an oomycete plant pathogen found in our data (Peronosporaceae). This suggests that the Zeale primers could potentially be used to identify mildew infections in apple orchards due to conserved regions shared between insects and oomycetes (Marquina et al., 2019; Rasmussen et al., 2021b). The analyses of metabarcoding data can also be limited by issues regarding incomplete DNA reference databases used for taxonomic identification (Deiner et al., 2017; Schenk et al., 2019; Rasmussen et al., 2021b). This is shown in our study where sequences were inaccurately assigned to species that were not previously documented in Danish apple orchards (*Lonchoptera uniseta* and *Periplaneta americana*) (Deiner et al., 2017; Ruppert et al., 2019; Schenk et al., 2019; Rasmussen et al., 2021b).

The study of pollinators in orchards till now has mainly been focused on Hymenoptera, which is considered the most effective pollinator (Fründ et al., 2013; Ramírez and Davenport, 2013; Dunn et al., 2020; Pardo and Borges, 2020; Rader et al., 2020). However, pollination in apple orchards includes a wider range of wild insects such as Coleoptera, Diptera or Lepidoptera (Rader et al., 2016; Lucas et al., 2018; Földesi et al., 2020). The findings from our visual census showed that, although non-Hymenoptera pollinators were abundant in apple orchards (47%), Hymenoptera was still the most abundant order observed (53%). This could be due to honey bee hives being placed within apple orchards to increase the success rate of pollination, explaining their high presence in our study (47%) (Corbet et al., 1991). Some challenges of using visual census for arthropod diversity assessment include long sampling time and expert knowledge in accurately identifying arthropods (Evans and Kitson, 2020), and being dependent on the weather (Ramírez and Davenport, 2013). Additionally, some of the most important crop pollinators such as bumblebees, hoverflies and wild bees can be small, cryptic, fast, and difficult to identify through visual census (Corbet et al., 1991; Leonhardt et al., 2013; Orford et al., 2015; Dunn et al., 2020; Rader et al., 2020; Rasmussen et al., 2021a). This could lead to an underestimation of pollinator presence and diversity assessed using visual census (Rader et al., 2016; Lefebvre et al., 2017).

Arthropod richness did not differ significantly across orchards with either metabarcoding or visual census. Some differences did emerge between orchards. The Pometum orchard flowered later than the other three orchards and was accordingly sampled about 10 days later, so the absence of some arthropod groups might reflect the later flowering. Secondly, interactions among flower visitors and suitable additional resources also play a role in shaping arthropod communities (Delaplane et al., 2000; Klein, 2011; Delpon et al., 2019; Valido et al., 2019; Rasmussen et al., 2021a; Weekers et al., 2022). High densities of honey bees have been demonstrated to detrimentally affect wild pollinator populations in the orchards as an effect of competitive exclusion (Klein, 2011; Valido et al., 2019; Angelella et al., 2021; Rasmussen et al., 2021a). This is also reflected in our study where we observed lower wild pollinators densities in the two orchards with honey bee hives placed inside the orchard (Frydenlund and Kildebrønde). Ventegodtgaard had the highest, though not significantly so, arthropod richness according to both methodologies as compared to Frydenlund and Kildebrønde. Hence organic management practice may have resulted in a higher arthropod presence in Ventegodtgaard (Gomiero et al., 2011; Ahrenfeldt et al., 2019; Samnegård et al., 2019).

### Comparison between metabarcoding and visual census

All arthropod orders found with both methodologies were previously recorded in apple orchards (Garcés and Soto, 2000; Rader et al., 2020). Coleoptera, Diptera and Lepidoptera were common across both methodologies. While metabarcoding found DNA traces from herbivorous Lepidopterans, which are pests of apple, and which by the time of sampling were in the larval stage, visual census showed few Lepidopteran species visiting the flowers. Apple orchards represent one of the most diverse cultivated agroecosystems with over 1000 arthropod species recorded (Szentkiralyi and Kozar, 1991; Šťastná and Psota, 2013). While pests are well known and described (Blommers, 1994), less is known about the communities of natural enemies and pollinators and how to augment them (Gomiero et al., 2011; Herz et al., 2019; Samnegård et al., 2019). The flowering season in Denmark is characterized by variable spring conditions, often windy and cold, which negatively affect arthropod activity (Ramírez and Davenport, 2013; Gibbs et al., 2017). While we selected sampling dates when minimum criteria for weather were met, still, a larger sampling effort may be needed this far north to detect certain groups such as hoverflies (Syrphidae) or wild bees (Apidae), which are more active in warm, sunny weather (Corbet et al., 1991; Fründ et al., 2013; Ball and Morris, 2015; Orford et al., 2015; Pardo and Borges, 2020). Here, metabarcoding can be extremely useful and allow us to study arthropod flower visitors’ presence regardless of foraging periods. The arthropods identified using metabarcoding and visual census included arthropods with different roles in the ecosystem (pollinator, pest, predator), both nocturnal and diurnal. Metabarcoding does not distinguish between developmental stages, and the Lepidopteran species detected from apple flowers by the method occurred in the larval stage at the time of sampling. This is demonstrated by some flowers that were seen with larvae and frass when sampling took place. In contrast, visual census data are derived from observing only adults which are considered as pollinators (CFIA, 2006; Turner et al., 2010; Deiner et al., 2017; Taylor, 2019; Rader et al., 2020).

Metabarcoding can provide species-level taxonomic identification from a wide array of arthropod diversity compared with traditional approaches (Ruppert et al., 2019; Evans and Kitson, 2020). Metabarcoding can also be a useful tool to study nocturnal or cryptic flower visitors that can be difficult to observe using visual census (Barnes and Turner, 2016; Deiner et al., 2017; Ruppert et al., 2019). Examples are Diptera and Lepidoptera which might be present in the flowers early in the morning or during the night (Elberling and Olesen, 1999; Ssymank et al., 2008; Rader et al., 2016; Nunes-Silva et al., 2020; Rader et al., 2020). Thus, the higher presence of Lepidoptera (37-49%) and Diptera (33-44%) detected with metabarcoding might be due to DNA traces being more recent and less degraded than other groups of arthropods when sampled in the morning (CFIA, 2006; Woodcock et al., 2014), or in the case of Lepidopteran larval traces, could be due to frass left on flowers (Feinstein, 2004).

Issues such as primer biases and incomplete DNA reference database used for taxonomic assignment probably led to the underrepresentation of typical pollinator taxa such as Hymenoptera and Syrphidae (Lamb et al., 2019; Marquina et al., 2019; Rasmussen et al., 2021b). Metabarcoding is very sensitive and may detect DNA traces transported from other locations by means of air or other arthropod flower visitors (Deiner et al., 2017; Lamb et al., 2019; Todd et al., 2020), which may in turn lead to misrepresentation of flower visitor diversity (Barnes and Turner, 2016). Visual census can provide a priori knowledge of the specific community being assessed, making it a valuable complementary tool to be used in conjunction with metabarcoding (Ruppert et al., 2019), also providing quantitative data. Even though metabarcoding reads can be converted into presence-absence data or relative read abundance (RRA), quantitative inferences of biomass are challenging as it depends on several well-documented variables. These include the starting DNA yield (larger species generate more reads), sample size (appropriate sample size to represent the sampled community) and primer selection (different primers can amplify different taxa within the same sample) (Blanckenhorn et al., 2016; Deagle et al., 2019; Lamb et al., 2019). Visual census on the other hand, can be biased by risk of double-counting and is dependent on the expertise of the observer, whose presence could lead to an alteration of arthropod behaviours (Deiner et al., 2017; Todd et al., 2020). Hence, the utilisation of either technique on its own has its limitations for arthropod diversity assessment and forthcoming studies could consider employing both in unison to get a better overview. In future studies, the metabarcoding workflow can be tweaked to optimise arthropod detection on flowers, such as using two or more primers, which will allow for adequate amplification of low-quality template DNA (Zhan et al., 2014; Ficetola et al., 2015; Marquina et al., 2019; Thomsen and Sigsgaard, 2019). This is consistent with other studies that support the need for multi-primer approaches to assess arthropod communities (Marquina et al., 2019). Increasing the number of biological and technical replicates, albeit at an increased cost and time, can also help to better detect rare and cryptic species (Deiner et al., 2017; Ruppert et al., 2019). DNA traces in flowers may be short-lived and inclusion of additional sampling in the afternoons may better capture traces of day-active flower visitors.

### Implications on orchard managements

Knowledge on beneficial and detrimental arthropods visitors to apple orchards is crucial to improving management techniques (Hambäck et al., 2021). Our study detected the presence of two important apple pests, *Operophtera* and *Yponomeuta* moths, that feed on developing apple fruitlets and leaves which can result in high economical losses if not controlled (Svensson et al., 1999; CFIA, 2006; Turner et al., 2010). Within the orders Diptera and Coleoptera, there are both apple pests and natural enemies. Adults from these two orders can also be pollinators (Levesque and Burger, 1982; Free, 1993; Compton and Key, 2000; Alford et al., 2003; Brown et al., 2007; Clement et al., 2007; Kazachkova, 2007; Steenberg and Harding, 2009; Orford et al., 2015; Kolcsár et al., 2016; Ouvrard et al., 2016; Rader et al., 2016; Thomsen and Sigsgaard, 2019; McKerchar et al., 2020; Pardo and Borges, 2020; Rader et al., 2020; Osterman et al., 2021). Arthropod predators have been used in apple orchards for pest control to reduce the use of insecticides and maintain apple production, yield, and quality (Porcel et al., 2018; Herz et al., 2019; Dunn et al., 2020; Hambäck et al., 2021). Results from this study confirmed the presence of the invasive ladybird *Harmonia axyridis*, a natural enemy of aphids and coccids, also established in Denmark (Steenberg and Harding, 2009). Additional resources are needed to ensure the presence of not only pollinators but also predators in managed ecosystems, such as apple orchards (Brown et al., 2007; Joshi et al., 2016; Saunders et al., 2016). Thus, data included in this study can allow farmers to select more efficient and specific pest control techniques while improving the surrounding landscape to attract certain beneficial arthropods that they know are visiting their orchards.

### Future studies

Agricultural systems are facing several threats such as pollinator decline and climate change (García and Miñarro, 2014; McKerchar et al., 2020; Pardo and Borges, 2020). Having a high diversity of arthropods in these systems is seen as a way to overcome some of these threats (Russo et al., 2015; Dunn et al., 2020; Hambäck et al., 2021). Hence, the study of flower visitors is an important step towards preserving agricultural systems (Klein et al., 2007; Leonhardt et al., 2013; Porcel et al., 2018).

We demonstrated in our study using both visual census and metabarcoding the existence and importance of wild non-bee pollinators such as Diptera, whose presence in apple orchards has been underestimated or even understudied in previous studies (Delaplane et al., 2000; Ramírez and Davenport, 2013; Rader et al., 2020). Therefore, management practice should consider the trade-offs of some arthropods with dual roles in orchards depending on their life cycle. Interactions across groups of arthropod flower visitors, as well as nocturnal and diurnal ones, should also be taken into consideration for future management practices. Awareness about possible complementarity and competition among arthropods could be necessary to improve fruit yield and quality.

Management practices (IPM, organic), orchard arrangement (number of rows, field size), additional floral resources (heterogeneous landscape, apple varieties) and interactions among arthropods (density and location of honey bee hives) can affect arthropod communities within the orchards (Delaplane et al., 2000; Leonhardt et al., 2013; Russo et al., 2015; Campbell et al., 2017; Földesi et al., 2020; Pardo and Borges, 2020; Rasmussen et al., 2021a). Future studies should include vegetation surveys of the landscape surrounding the orchards, as well as addressing the possible influence of management variables, such as orchard arrangement, on arthropod activity.

To conclude, our study provides the starting point for a more complete overview of biodiversity of arthropod flower-visitors found in apple orchards. The combination of molecular and traditional non-molecular techniques for arthropod assessment is complementary and can overcome some of the limitations inherent with using only one method. Going forward, we recommend the use of both molecular and non-molecular approaches in the assessment of arthropod diversity if time and budget permit. Otherwise, utilising a molecular approach such as metabarcoding with at least two primers can help to optimise arthropod detection. The outcomes of our study can support future management practices moving towards more resilient and environmentally friendly agricultural systems.

## ACKNOWLEDGMENTS

Research by LS and NGG was funded in part by the BEESPOKE project “Benefitting Ecosystems through Evaluation of food Supplies for Pollination to Open up Knowledge for End users” J-No.: 38-2-2-19, founded by the Interreg VB North Sea Region programme. PYSC was supported by the European Union Horizon 2020 research and innovation programme under grant agreement No 765000, H2020 MSCA-ITN-ETN Plant.ID network. For generating the Illumina data, we would like to thank the staff at AllGenetics. We would like to extend our gratitude to Karen Jensen and Helle Mathiasen (SOBI) for helping us to collect samples in the field. Lastly, we appreciate the help that Pablo Castro, David Escobar and Ismael Rodriguez who provided for the biodiversity and statistical analyses.

## SUPPLEMENTAL INFORMATION

- README-This file contains a summary of all the supplementary material contained within this folder.
- Supp1-Supplementary Tables S1 to S13
- Supp2-Supplementary 2-Figures S1 to S2
- rawdata.tgz-Raw sequence files from AllGenetics
- cutadapt_OBItools.zip: Cutadapt trimmed files for OBITool
- My_code_pollinators.bash: Bioinformatic script used to generate metabarcoding data contained in Supp1
- paper_Rscript-R scripts used to generate metabarcoding and visual census data (Rmd) contained in Supp1 and Supp 2
- paper_Rscript-R scripts used to generate metabarcoding and visual census data (html) contained in Supp1 and Supp 2
- paper_Rscript.zip-.csv files used for stats test on R-studio

## AUTHOR CONTRIBUTIONS

NGG, PYSC and LS designed the research; NGG and LS collected samples; NGG, DHS and AllGenetics performed laboratory work; NGG and PYSC did the bioinformatic analysis; NGG did the statical analysis; NGG made the figures and wrote the paper with input from PYSC and LS. All authors contributed to the final version of the submitted manuscript.

## DATA RESOURCES

Raw Illumina sequencing files, trimmed files used for analyses, R and bioinformatics scripts used for data generation and .csv files used for statistical analyses are available at the University of Copenhagen Electronic Research Data Archive (UCPH ERDA) Digital Repository, https://doi.org/10.17894/ucph.12092ac6-80c4-4c8c-9572-208981c0c5e2, and will be made available on the DRYAD Digital Repository upon acceptance of this manuscript.

