## Supplemental 2- Figures S1 to S2 for "Assessing flower-visiting arthropod diversity in apple orchards through environmental DNA flower metabarcoding and visual census"

**Table of Contents:**

| **Fig. S1 Metabarcoding rarefaction analysis** | Page 2 |
| --- | --- |
| **Fig. S2 Visual census rarefaction analysis** | Page 3 |

### S1. Metabarcoding rarefaction analysis

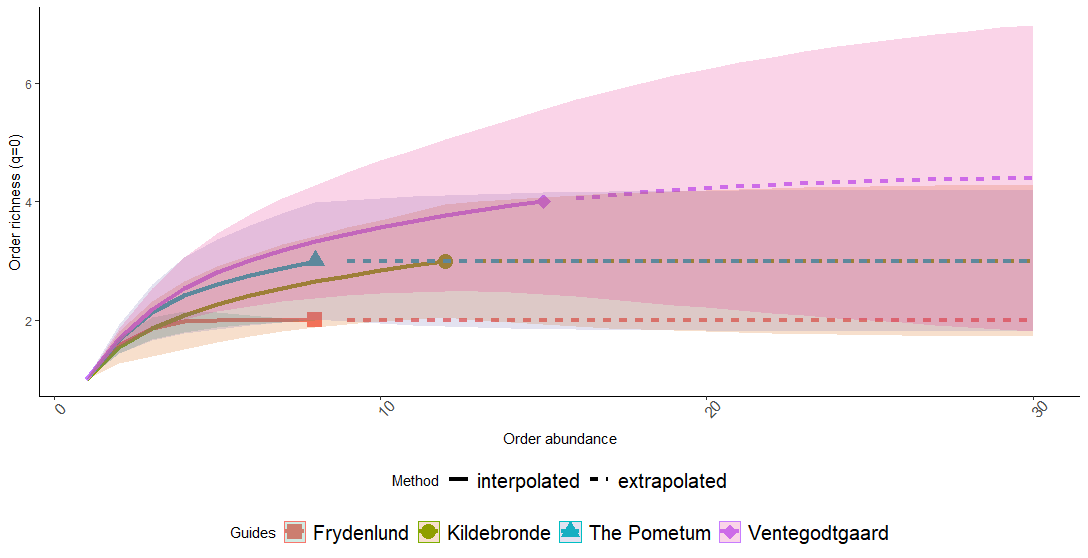

**Figure S1.** Order abundance-based rarefaction (solid) and extrapolation (broken) curves for order richness (q=0) of arthropods visiting apple orchards based on metabarcoding. 95% confidence intervals (shaded areas) obtained by bootstrap method based on 30 replications.

### S2. Visual census rarefaction analysis

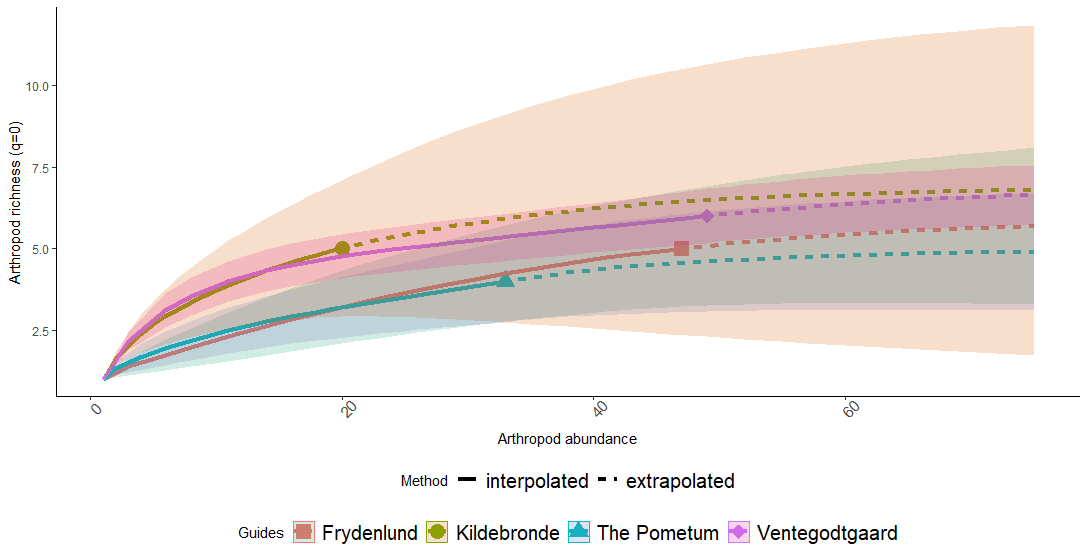

**Figure S2** Arthropod abundance-based rarefaction (solid) and extrapolation (broken) curves for morpho-group richness (q=0) in apple orchards at all sites based on visual census. 95% confidence intervals (shaded areas) obtained by the bootstrap method based on 75 replications.
